# Mechanistic classification of the AAA superfamily with protein language models

**DOI:** 10.64898/2026.10.05.756829

**Authors:** Jeffrey A Swan, Christopher P Hill

## Abstract

The **<u>A</u>**TPases **<u>A</u>**ssociated with various cellular **<u>A</u>**ctivites (AAA) are a class of proteins with diverse structure-function relationships whereby conserved 3-dimensional architecture is employed in varied mechanistic contexts. While cryo-electron microscopy has revealed extensive mechanistic data, predicting how specific sequence elements dictate unique biochemical mechanisms remains a challenge. Here, we present a machine learning framework to resolve these sequence-function relationships. We established a hierarchical classification scheme based on five distinct modes of coupling ATP hydrolysis to mechanical work. Using **CatFunc**, a lightweight protein language model (PLM)-based pipeline, we leveraged evolutionary context from millions of protein sequences for supervised classification of AAA domains based on mechanism. We observed excellent recall of our mechanistic annotations in masked validation studies, demonstrating that domain level context is sufficient to encapsulate the various observed mechanisms. Notably, systematic misclassifications revealed a shared evolutionary and mechanistic ancestry within the rotary motor class. Applying the model to out-of-distribution proteomic datasets, we established metrics to detect unrepresented functions. Our framework provides a scalable strategy for decoding molecular function from sequence data, offering a transferable approach for other biomedically important families such as membrane transporters, G protein-coupled receptors and kinases.

## Introduction

### The AAA superfamily of ATPases

**<u>A</u>**TPases **<u>A</u>**ssociated with various cellular **<u>A</u>**ctivites (AAA) comprise one of the best studied and functionally diverse family of protein coding genes (*1–3*). Members of this superfamily are defined by the presence of one or more distinct ATPase sequences, called AAA domains, within their open reading frame (ORF). The AAA domain is comprised of a distinct pattern of secondary structures (**Fig. 1A**) with conserved Walker A and Walker B motifs (*4*) at defined positions where they effectuate ATP binding and hydrolysis, respectively. AAA domains exhibit a conserved 3D structure (**Fig. 1B**) composed of large and small domains that can assemble into higher order oligomers with ATP molecules situated at the protomer interfaces (*5*). Interactions at these interfaces are intertwined with the enzymatic cycle of nucleotide binding, hydrolysis and release, leading to chemo-mechanical coupling of ATPase activity with a molecular biological function (*6*). These properties make AAA-domain containing targets ideal for structure-function studies using cryo-electron microscopy, as the target complexes are ideal size for the technique, which can resolve the γ-phosphate of ATP, and thereby distinguish coexisting pre- and post-ATP hydrolysis structures within heterogeneous samples. This has led to a wealth of structural and mechanistic information for this class of ATPases. For reviews on the mechanistic insights on AAA proteins see references (*7–9*).

**Figure 1:**
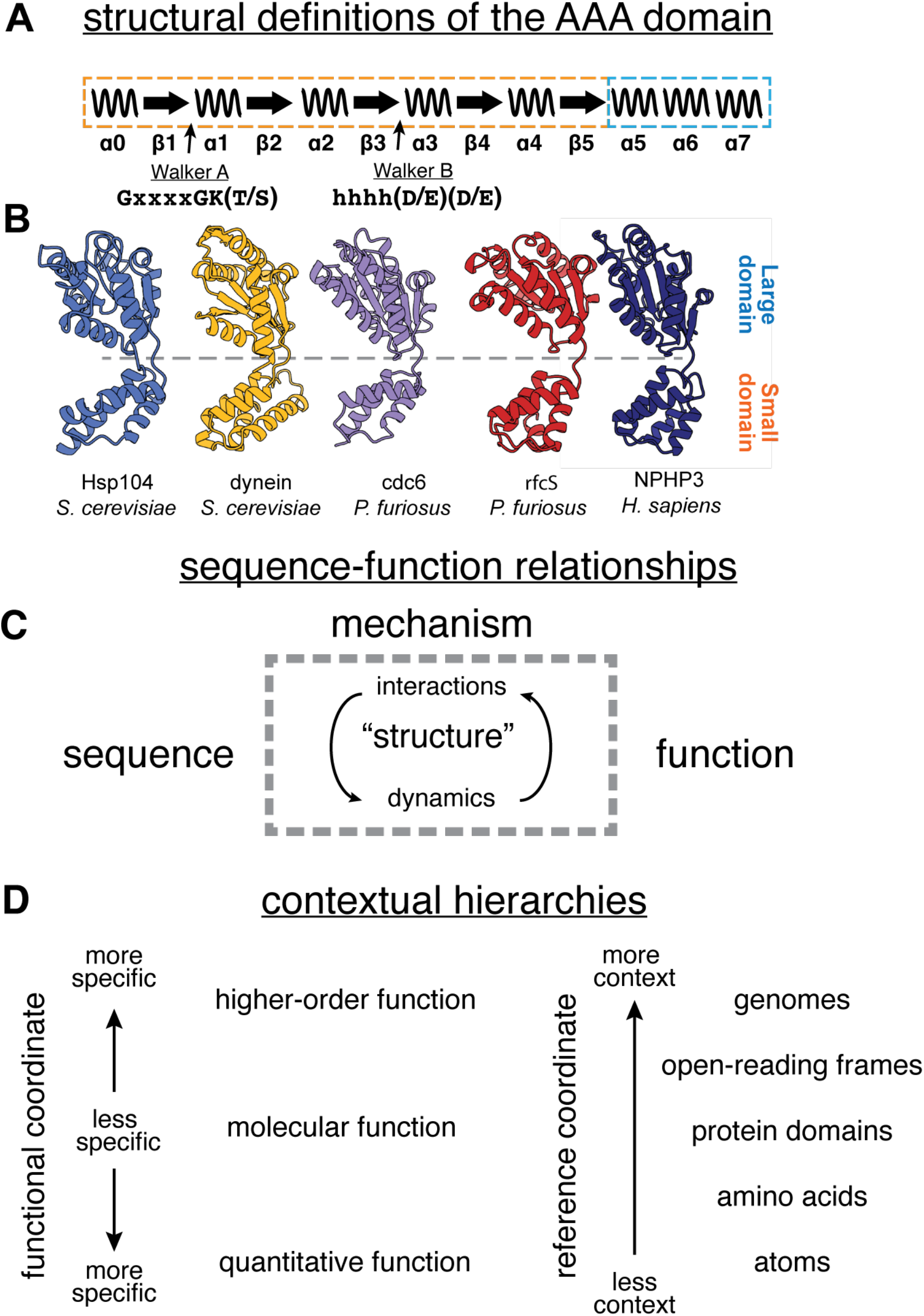
Sequence function relationships in the AAA superfamily. **A**) Secondary and primary structural definitions of the AAA domain. The AAA domains is defined by a specific pattern of alpha helices (waves) and beta sheets (arrows), with Walker A and Walker B motifs at set positions within this secondary structural pattern. Orange and blue dotted lines denote the large and small domains. **B**) 3D structures of AAA domains from five mechanistically diverse proteins. **C**) Schematic representation of the concepts connecting protein sequence to function. Sequence and function are rarely, if ever, connected by a static 3D structure, but rather involve a complex dance of structural dynamics and interactions with other factors such as substrates or regulators. We use the term mechanism to encompass these factors broadly. **D**) Contextual reference frames for functional inference. Biological function inherently occupies various frames of reference. To ensure that the necessary information for inference is encompassed within the task, the scope of input information must be matched to scope of biological function.

While AAA domains generally possess a conserved mechanism for ATP hydrolysis, whereby a catalytic magnesium ion is coordinated by acidic residues in the Walker B motif (*10*), the functional consequence of this catalytic event can be utilized in a processive or non-processive manner, corresponding to a motor function observed in protein unfolding and subcellular locomotion, vs. regulatory functions observed in DNA replication and signal transduction (*11*). Even within the motor class, distinct mechanisms were proposed early on, distinguishing AAA variants that project force through the central pore from those that exert torque around the outside of the ATPase ring (*12*). This leads to the central hypothesis of this work: that molecular mechanisms are encoded locally within the AAA domain level, rather than exclusively by contextual factors such as N or C-terminal extensions or binding partners, leaving a reliable evolutionary signal within the AAA domain itself that can be used for inference of molecular function.

### Challenges in formalization of the sequence to function problem

Advances in machine-learning based approaches to structural biology, such as AlphaFold (*13*), have made it much simpler to obtain experimental quality structures. However, these approaches have limited utility in understanding the sequence-function relationship, since molecular function arises out of a complex dance of intermolecular interactions, chemical regocnition and catalysis and conformational changes (*14*) (**Fig. 1C**). Furthermore, function is an inherently contextual problem (**Fig. 1D**) that cannot necessarily be formalized to a single objective in the same way that sequence-based lowest-energy structure prediction can. For example, molecular functions such as kinase or ATPase activity may be encoded within a given protein domain, while their higher order biological function may be determined by external regulators that require additional context to predict accurately. To complicate matters further, biological function can be influenced heavily by quantitative aspects of function such as binding affinity or enzymatic activity, which arise from detailed physical properties of the biomolecules being considered. Attempts to predict such quantitative aspects of function the evolutionary level have not typically been successful (*15*) and will likely require atomistic models to predict from first principles.

### Protein Language Models as a tool for sequence-function analysis

Protein language models (PLMs) are emerging tools that treat protein sequences as a naturally occurring language (*16*) whose internal residual stream can be utilized for transfer learning tasks (*17*) related to protein sequence analysis and structure prediction (*18–23*). Implementations such as the Evolutionary Scale Model (ESM) (*24*) are transformer-based neural networks trained on the self-supervised masked language modeling task (**Fig. 2A**). Here, the language model is challenged to predict masked portions of protein sequence from a large corpus such as the UniREF50 database. To achieve this task, the model learns to predict evolutionary context of a given protein sequence, which are represented internally within weights of the trained language model. The language model is composed of multiple layers, through which these numerical representations are passed at inference time, the values associated with the nodes are known as a residual stream and can be conceptualized as high-dimension contextual embeddings of proteins sequence, referred to herein as embeddings. These embeddings can then be used as numerical representations of a protein’s evolutionary context (**Fig. 2B**), that can be used for various classification or regression tasks to make predictions about a protein sequence of interest (*23*).

**Figure 2:**
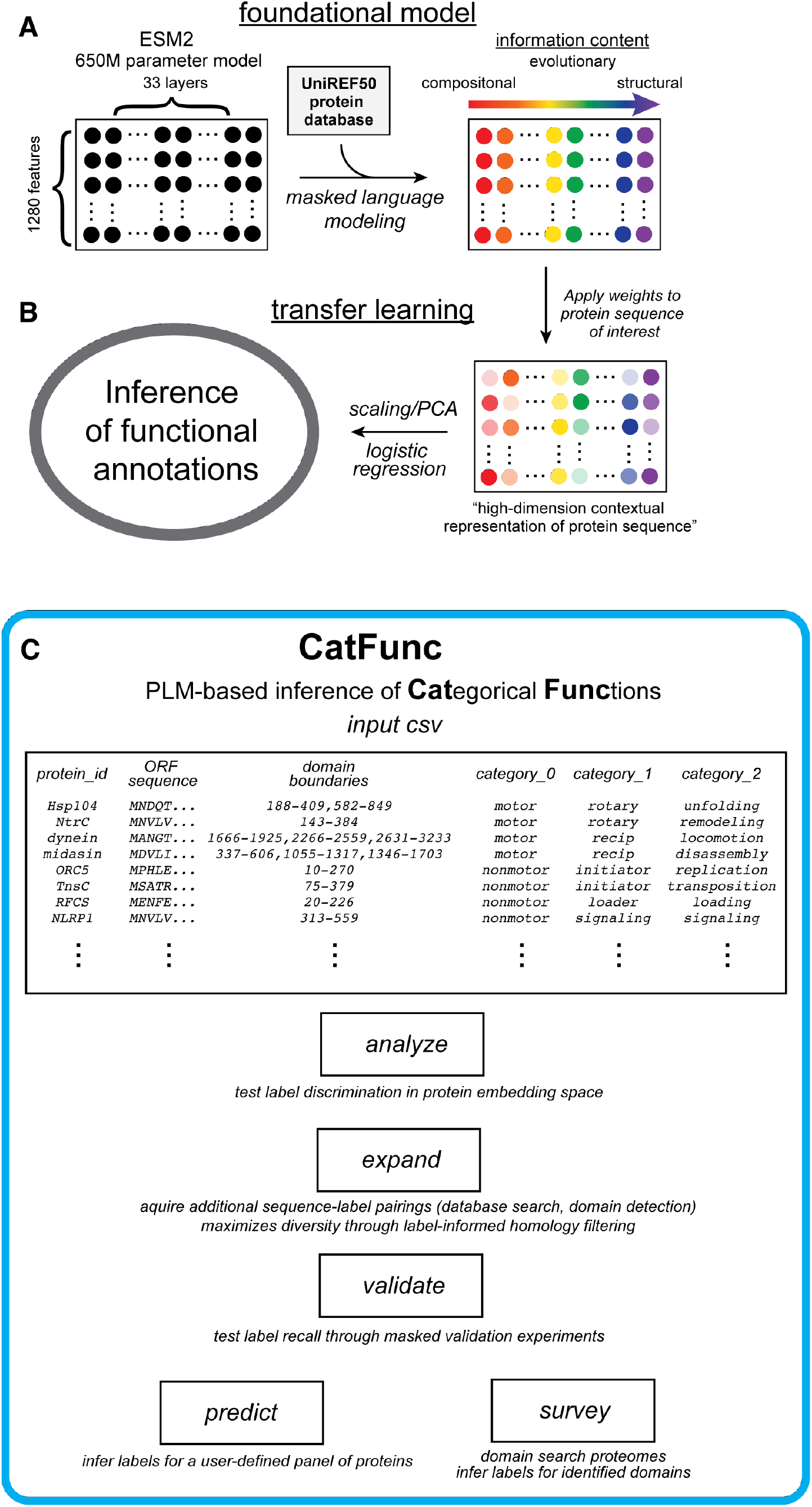
PLM-based transfer learning of user-defined annotations with CatFunc **A**) The ESM2 protein language model is pre-trained on masked-language modeling against the corpus of protein sequences observed in nature. Through its training, differen contextual logic is encoded in its neural network, with different types of information present throughout its layers. These pretrained weights can be applied to a protein of interest for transfer learning (**B**), resulting in a numerical representations of the proteins evolutionary context, often referred to as ‘embeddings’. In this study, we associate these embeddings with mechanistic annotations and use them for functional inference through supervised classification. **C**) The user provides an input file containing the protein sequences of interest, domain boundaries restricting the frame of reference for the protein sequences, and a list of categorical annotations. CatFunc the performs a series of modular analyses. The ‘analyze’ module does a cursory check of how well the annotations provided are discriminated in ESM2 embedding space across the various layers. The ‘expand’ module uses database searches to acquire additional protein sequences and annotations, and enriches diversity within the dataset by label informed homology filtering. The strength of the evolutionary signal within the expanded set of protein embeddings is tested using the ‘validate’ module. In the ‘predict’ module, annotations are generated user-defined list of protein sequences using the entire expanded dataset of annotated protein sequences for training. Proteomic datasets are scanned for the domain of interest, and annotations are generated based on the logistic regression model trained on the expanded dataset.

Early work on the ESM family of PLMs demonstrated that knowledge 3D-structure emerges within the final layers of the language model (*25*), in the form of attention between residues corresponding to evolutionary covariation relationships, with less specific evolutionary trends believed to be encoded in the earlier layers (*26, 27*) (**Fig. 2A**). Notably, the contact maps emerging from the final layers of the model also contain many “false positives,” (*24*) evolutionary contacts are also observed that do not correspond to 3D contacts in the structure of the protein. This observation underscores the idea that PLM-based embeddings as a structure agnostic tool function inference, using them as numerical representations of a protein’s evolutionary that can be subsequently used by traditional machine learning methods for various classification or regression tasks.

Here we leverage PLM-based representations of protein sequence against the wealth of diverse mechanistic data on AAA domain containing proteins to implement a systematic, structure-agnostic approach for inference of molecular function. Using this analysis as a foundational case study, we developed CatFunc (**Fig. 2C**), a computational pipeline whereby user-defined hierarchical categorical annotations can be assigned to protein sequences, and the evolutionary encoding of these annotations investigated and projected onto unknown examples using PLM-based inference powered by the ESM2 model. In the implementation of CatFunc, we begin by manually curating a foundational set of AAA domains with reliable mechanistic annotations and embed a method for expansion of this dataset to hundreds of examples, even with strict homology restraints enforced. The evolutionary signal associated with these mechanistic annotations within expanded dataset was tested using masked validation studies focused on global ESM embeddings of domain or full ORF sequences. This machine learning pipeline for functional inference and applied to AAA domains from either a user-defined list of AAA domains with uncharacterized mechanisms, or AAA domains extracted from various proteomes across the kingdoms of life, providing metrics to identify outliers from the originally proposed mechanistic classes. Together CatFunc encompasses a general schema for inference-based structure/mechanism studies and establish a sequence-based approach to identify novel molecular functions for experimental interrogation.

## Results

### Hierarchical mechanistic classification of AAA proteins

We hypothesized that the differences between the mechanisms of different AAA proteins are encoded within the isolated AAA domain, and that common sequence patterns manifest these mechanistic features. As an initial step to test this idea, we did an extensive literature search to track mechanistic diversity. Through this extensive review, we observed sufficient mechanistic data to assign eight distinct molecular functions to groups of AAA-containing proteins (**Fig. 3A, Extended Data Table 1, Supplementary Data 1**), as well as several AAA proteins that did not fit neatly into these categories, or for which insufficient data was available to assign labels. The initial set of 122 labeled AAA domains across 103 ORFs with mechanistic labels is referred to herein as the literature set. We then grouped these eight major archetypes into five distinct mechanisms with respect to how ATP-hydrolysis is coupled to mechanical function: rotary motors, reciprocating motors, initiators, loaders and signal transduction ATPases. Two even coarser groups emerged from our mechanistic labels, which we named the motor and non-motor classes, referring to the functional output of chemo-mechanical coupling.

**Figure 3:**
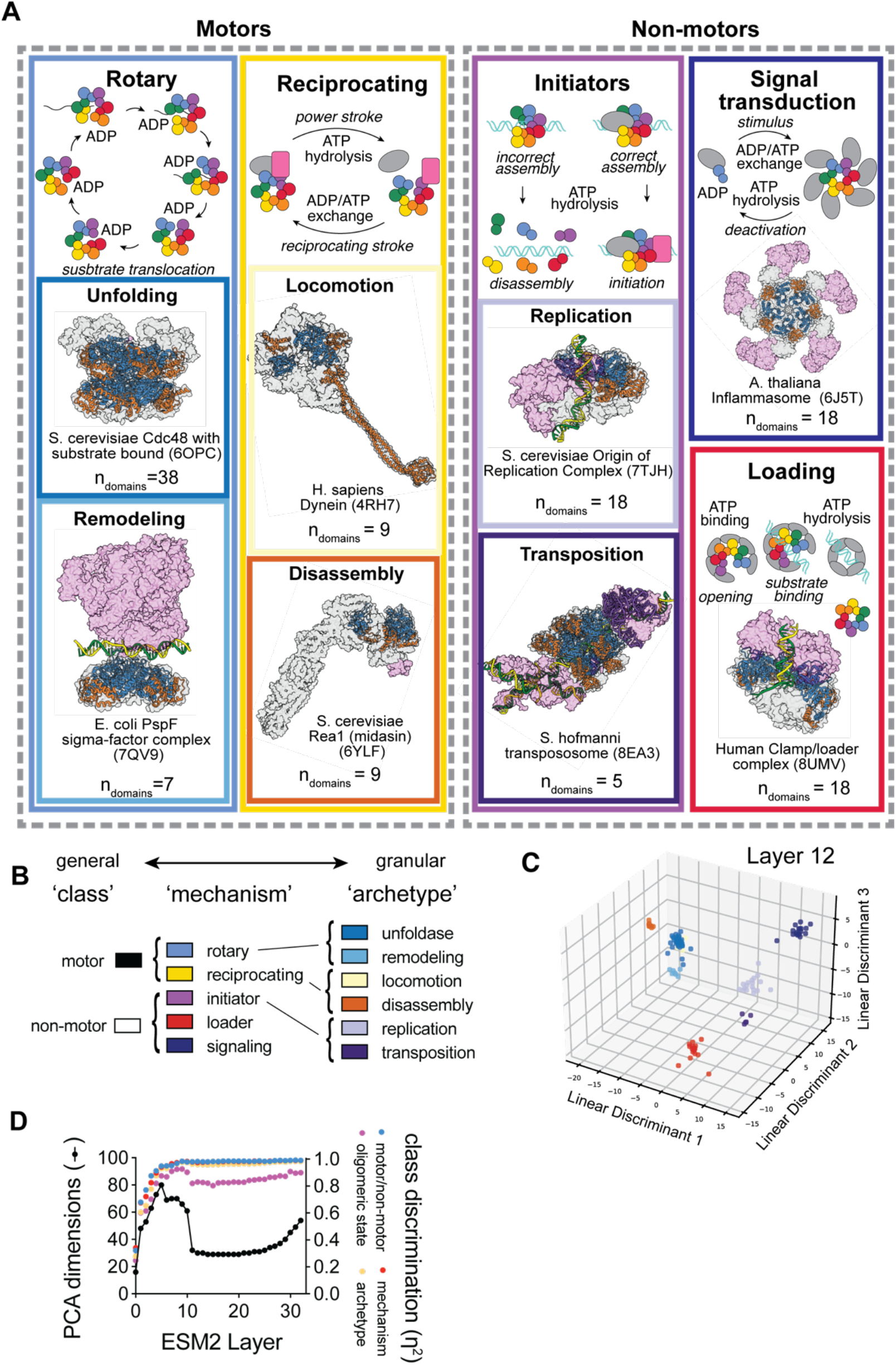
Literature search and embedding analysis of hierarchical mechanistic classes of AAA proteins. **A**) Various examples from a literature search of AAA domain containing proteins with experimentally determined mechanisms. For each mechanistic class, cartoons represent the experimentally determined common mechanism. Structural data are represented for example from each archetype, with large and small AAA domains shown in blue and orange, respectively. Protein structures from within the same ORF as the AAA domain are depicted as gray surfaces, and accessory factors are depicted as pink surfaces. **B**) ESM2 embeddings are depicted in 3-dimensional linear discriminant space for the 8 archetypes identified in the literature search. Archetype annotations are mapped onto each protein using the color coding defined in panel **A**. **C**) Per-layer principal component analysis of protein embeddings from the 122 AAA domains identified in the literature search. Per-layer analysis of the linear discriminants for each of 4 categorical annotations within the same set of protein embeddings are overlaid.

This hierarchical mechanistic classification system (**Fig. 3C**) comprised the inputs for our implementation of the general tool, CatFunc, that we developed to formalize and reproduce this analysis for other protein families. The next step in this implementation was to test how well distinguished these labels are amongst the literature AAA protein sequences in ESM2 embedding space. We therefore projected the foundational AAA domain sequences into Linear Discriminant (LD) space. In this analysis, the dimensions of the embeddings were reduced while keeping the projection that best discriminate the labels associated with each of our mechanistic categories. Figures **3B** and **S1** depicts ESM2 embeddings from the literature dataset projected into 3-dimensional space, optimized for linear discrimination of the ‘archteype’ labels, which are mapped onto each domain’s embedding using the colors defined in **3A**. Notably, in addition to each class clustering visually, both classes within the ‘rotary motor’ and ‘initiator’ mechanisms cluster together, despite the hierarchical information not being included in the LD analysis.

To quantify the discrimination at each layer of the ESM2 embeddings, we calculated η^2^, a measure how well separated the labels from each hierarchical category are in LD space with n_labels_ -1 dimensions for each category (**Fig. 3C**). This analysis showed that while discrimination was generally poor in the early layers, good separation was obtained by around layer 10. Consistent with this, we found that Principal Component Analysis (PCA) required more dimensions to represent 95% of the variance in the data in the early layers, while this number dropped precipitously around the same layers where LD of the labels in the literature embeddings rose. this suggests that the compositional information present in the early embedding layers acts evolutionary noise that is reduced in intermediate layers of the ESM2 embeddings, consistent with the idea that these middle layers converge upon a common evolutionary ancestry present within our structurally conserved dataset.

For comparison, we included a binary annotation encompassing whether each AAA domain acts within the context of heteromeric or homomeric oligomers as their functional unit. This label is asymmetrically distributed across the dataset (**Extended Data Figure 2**), with homomeric complexes only observed in the unfoldase, remodeler, transposition initiator and signaling ATPase archetypes. When subjected to the same per-layer LD analysis, this label exhibited consistently lower η^2^ values, indicating that it is not as strongly encoded in the global ESM2 embeddings as our mechanistic labels. This provides an important point of comparison to demonstrate the strength of the evolutionary signal underlying our mechanistic labels.

### Dataset expansion through contextual inference

Our initial literature search yielded 103 AAA domain-containing proteins for which we could confidently assign mechanism based on experimental data, with 122 individual AAA domains contained within those ORFs (**Extended Data Table 1**, **Fig. 4A**). We sought to expand this number using database searches, however, assignment of our mechanistic labels could not be done by simply searching against the domain sequences themselves, since it is unknown whether these mechanistic functional properties are encoded within the AAA domain, or whether additional context required to define them. To overcome this limitation, we designed CatFunc to perfrom Basic Local Alignment Search Tool (BLAST) searches using the full ORF of each AAA protein (**Fig. 4A**), and hits were filtered such that at least 85% of the hit aligned with the query sequence (see **Supplementary Data S3** for filtering statistics). Annotations were then transferred to the hits arising from each query.

**Figure 4:**
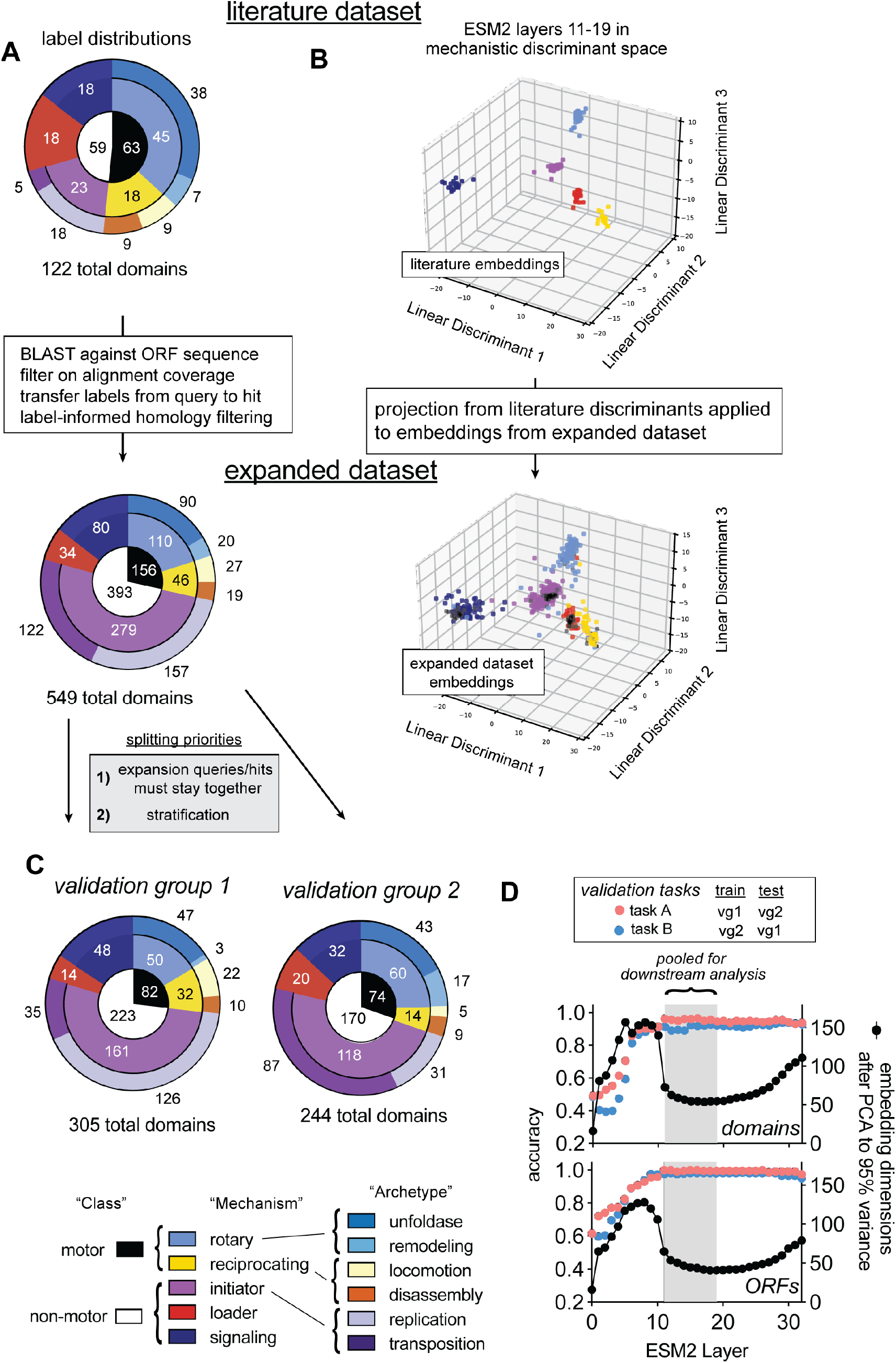
Dataset expansion and model validation for mechanistic classification of AAA domains. **A**) Distributions of the various hierarchical annotations of AAA domains within the foundational (literature) and expanded datasets depicted as concentric pie charts with the coloring scheme used throughout this manuscript to denote the labels within each category. **B**) Pooled literature embeddings from layers 11-19 projected into linear discriminant space for mechanistic annotation, with the mechanistic annotations mapped onto each embedding as color. Below, the projection that was derived for linear discrimination of the mechanistic annotations within the literature set is applied to the expanded dataset, with the literature embeddings depicted in black, and mechanistic annotations mapped onto the domains obtained through dataset expansion. **C**) Label distribution within the two groups of annotated proetein sequences, split from the expanded datset for validation. **D**) Per layer analysis of the expanded dataset’s principal components (connected black dots) and recall of masked annotations for each validation group by a logistic regression model trained on embedding from the opposite group.

AAA domains were extracted from this expanded ORF dataset using the hidden Markov model-based tool HMMer with a profile generated using AAA domains from our literature set, as well as some unclassified AAA domains (see **Supplementary Data 2** for the alignment used for domain boundary identification). To enrich diversity within this dataset, we then applied increasing stringent maximum pairwise homology filters (**Extended Data Figure 2A,** for the full expanded dataset before homology filtering see **Data S3**), while retaining sequences that would maximize inclusion of literature sequences and representation from each category of annotations. After applying the most stringent, 30%, homology cutoff we obtained a dataset of 549 AAA domains arising from 538 distinct ORFs, with double digit representation from the rarest labels (**Fig 4A**, **Supplementary Data 4**).

To assess the efficacy of our strategy for transfer of mechanistic annotations based on ORF homology, we analyzed the ORF sequences corresponding to our 30% homologous expanded domain set phylogenetically (**Extended Data Figure 3**). We then mapped mechanistic labels mapped onto the resulting phylogenetic tree to see how well they cluster. In general, archetype labels clustered near each other, suggesting that transfer of labels between these groups was appropriate based on the evolutionary signal encoded within their ORF structure.

As a qualitative test of whether the mechanistic labels of the domains within the expanded dataset could be reasonably discriminated on the same basis as the literature dataset, we took an LD projection for the ‘mechanism’ labels of the literature embeddings and applied it to embeddings from the expanded dataset (**Fig. 4B**). We observed reasonable separation of the mechanism labels in the expanded dataset domain embeddings on the same basis as the literature embeddings, suggesting that the same evolutionary trends in the literature dataset persist in the expanded dataset for these labels.

### PLM-based supervised mechanistic classification of AAA domains

Next we set out to validate a machine-learning pipeline designed to predict the hierarchical mechanistic annotations system for AAA domains. To do this, we designed CatFunc to split the expanded dataset into two groups (**Fig. 4C**) with BLAST queries grouped with their hits, and archetype labels stratified to the extent possible. CatFunc then applies a simple PLM-based machine learning pipeline (**Fig. 2A,B**) whereby logistic regression, a traditional method for supervised classification, was used to recall the mechanistic labels from each layer of our ESM2-derived representations of the expanded dataset (**Fig 4D**). Consistent with our LD results, we observed model performance maximize around layer 11, concomitant with a drop in the number of dimensions required to retain 95% variance in the data by PCA. This effect was observed for both ORFs and domains, with the ORF context window plateauing near 100%, with the domain level validation laggings lightly behind around 90%.

For a more comprehensive analysis of the mechanistic predictions, we next combined layers 11-19, where accuracy was highest, and applied the logistic regression model to domains and ORFs from the expanded dataset, this time tracking recall and Area Under the Receiver Operator Curve (AUROC) to gauge model performance (Figs 5A). Consistent with our initial LD analysis of the literature dataset, the model exhibited consistently higher recall values for our mechanistic labels than for oligomeric state (**Extended Data Figure 3A**), indicating that these annotations are better encompassed within the global ESM2 embeddings used for this study. The ‘archetype’ (n = 8) labels also showed lower recall on average (**Extended Data Figure 3B**). Examination of the per-class recall values for archetype labels (), shows that the average values are impacted by low performance amongst the classes that had very low representation in the training set for certain validation tasks (**Extended Data Figure 5A**, **Fig. 4C**).

**Figure 5:**
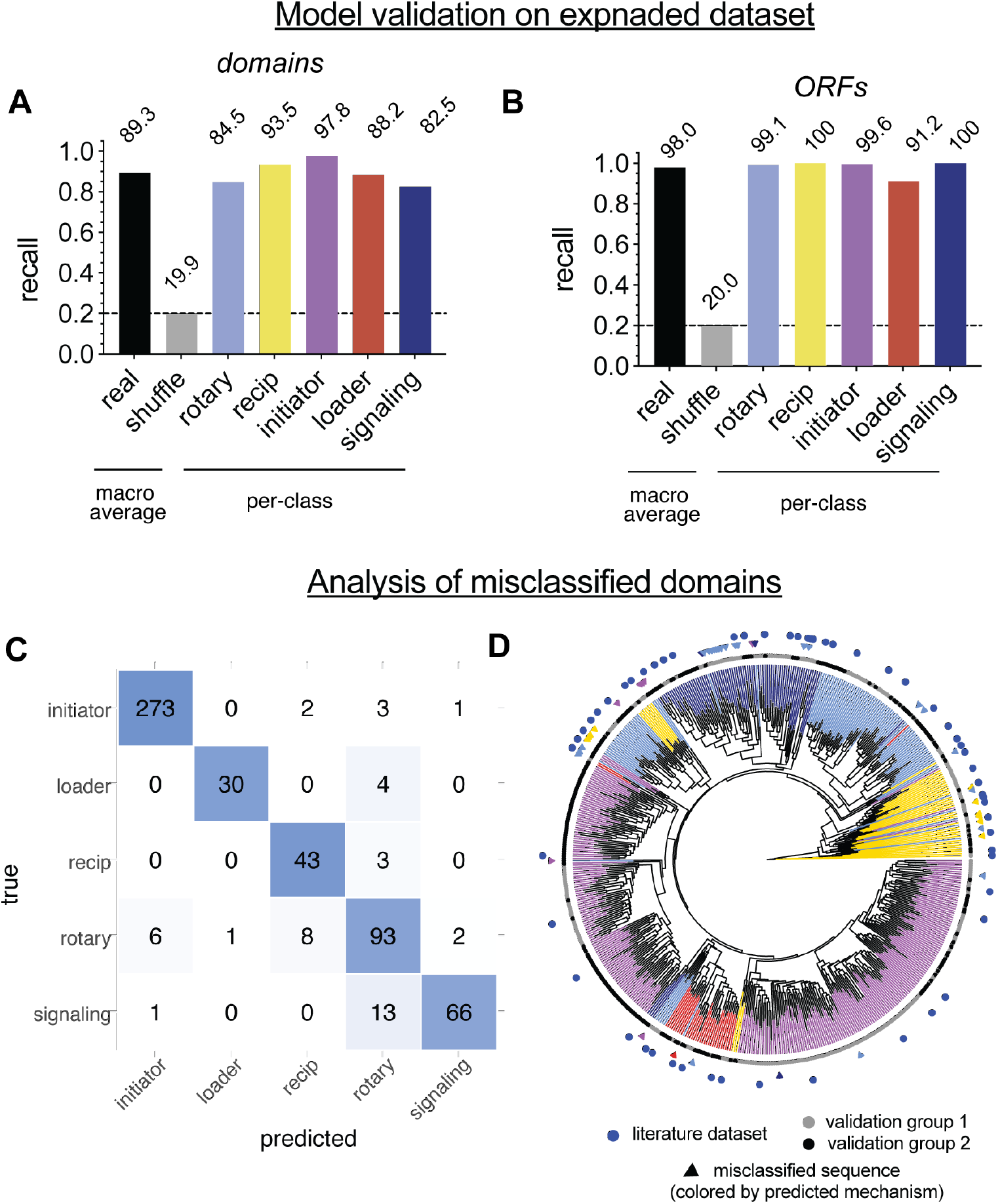
Internal validation of mechanistic inference within the expanded dataset of AAA domains. **A**) Macro-averaged and per-class recall of CatFunc’s validation module across both validation tasks, using logistic regression on ESM2 embedding layers 11-19 for AAA domain or **B**) full ORF sequences to recall mechanistic labels. **C**) Confusion matrix of domains. **D**) Phylogenetic anlayis of the expanded dataset of AAA domains, with literature values, validation groups and misclassified results mapped onto the tree. **E**) Confusion matrix of ORFs **F**) Phylogenetic analysis of ORFs from expanded dataset, with additional information mapped on as in (**D**).

Figures **5A,B** summarize the pooled layer validation studies, aggregated for the two validation tasks. Domain level prediction recalled almost 90% of the mechanistic labels, while including the full ORF context resulted in average recall of about 98%, suggesting that the additional context was useful for drawing functional correlations amongst the global protein embeddings. Within the domain-level predictions, the signaling label had the lowest per-class recall at around 83%. Misclassification was far less common in the ORF-level validation data, however the most common case was loaders or rotary motors being misclassified as initiators (**Extended Data Figure 3**). These were primarily misclassified as rotary motors (**Fig. 5C**, **Supplementary Data 4**), and clustered together phylogenetically (**Fig. 5D**) near signaling ATPase domains. The second most misclassified mechanistic class was the rotary motors themselves, which were often called as initiators or reciprocating motors (**Fig 5C**). The misclassified domains were often found as phylogenetic outliers, i.e. their placement in the domain-level phylogenetic analysis was often surrounded by other mechanistic labels, likely reflecting the fact that phylogenetic and PLM-based analyses call on the same trends to cluster the protein sequences.

### Out of distribution mechanistic inference on AAA domains

With our validated model for mechanistic inference trained on a relatively small subset of well-studied examples, we next sought to infer mechanisms from a more comprehensive set of naturally occurring AAA domains. CatFunc has two separate modules that can perform this function. First, categorical labels can be inferred for a user-defined panel of domains. **Extended Data Table 2** contains a panel of AAA domains for which we could not confidently assign mechanistic labels as well as CatFunc’s predictions on these (see **Supplementary Data 5** for predictions).

Additionally, we included a module within CatFunc that identifies novel domains of interest from proteomic datasets. To test this and critically examine the outputs, we used the same foundational alignment of AAA domains that was used to identify domain boundaries (**Supplementary Data 2**) during dataset expansion and used it to scan various proteomes (**Extended Data Table 3**) for novel AAA domains (**Data S6**). Notably, the proteome scans turned up several hits that contained annotations for ABC-transporters, a class of proteins involved in ATP-dependent translocation of substrates across the cell membrane, thus hits from our scan with these functional annotations were manually removed from the analysis (**Supplementary Data 7**).

**Figure 6A** summarizes the predicted abundance of our five mechanistic annotations in the AAA domains extracted from proteomes after removing the hits containing annotations as ABC-transporters. Rotary motors were the most abundant class across the genomes tested. Reciprocating motors were observed almost exclusively in eukaryotes, with only a small number of exceptions in E. coli. Initiators, despite being the most abundant class in the training data, was observed at consistently lower frequency across all the proteomes tested.

**Figure 6:**
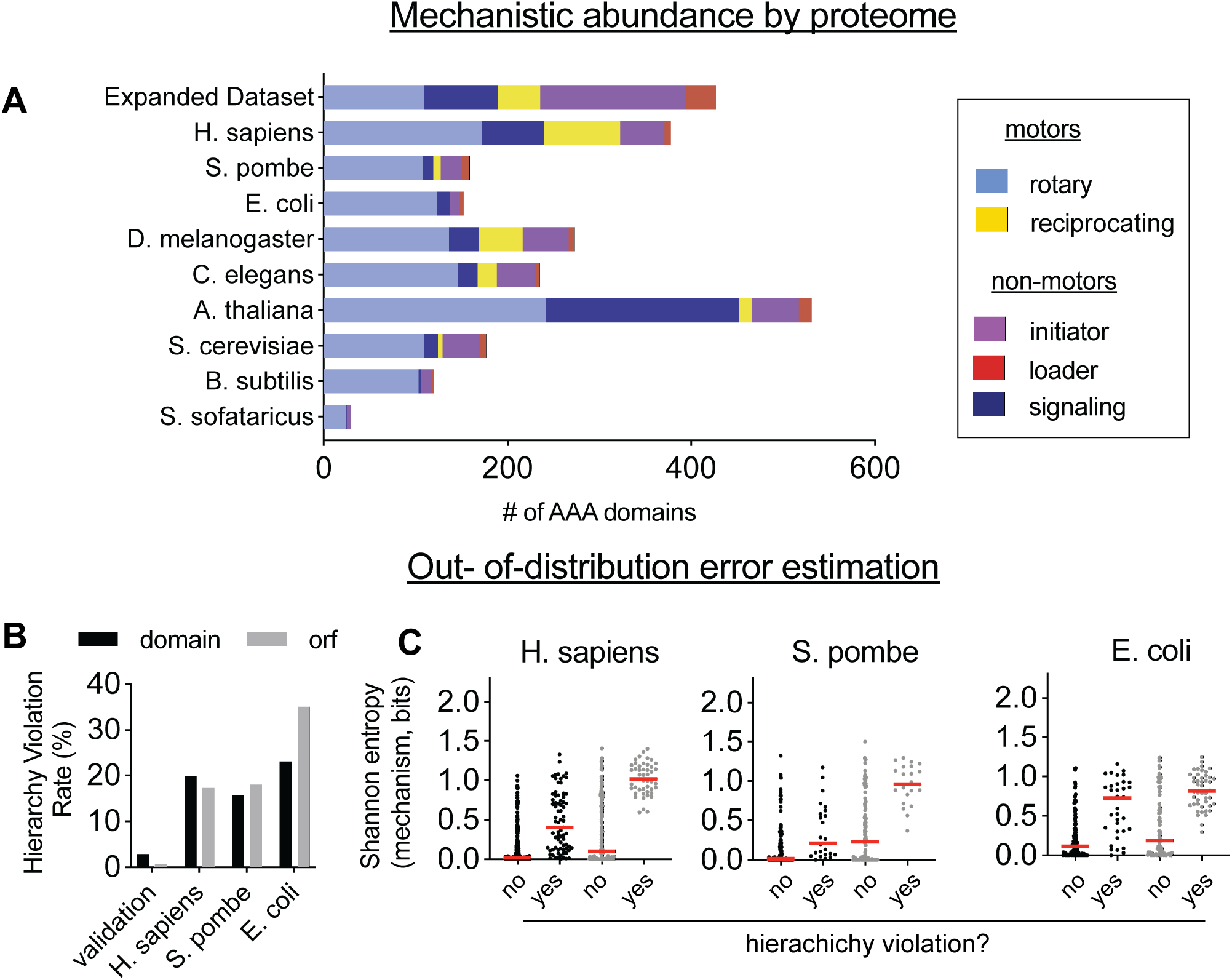
Mechanistic inference on proteomic AAA domains. **A**) Distribution of mechanistic classes within expanded dataset and AAA domain datasets derived from various proteomes. **B**) Prevalence of predictions that violate the hierarchical categories for AAA domain mechanism in a subset of the proteomes tested. **C**) Comparison of Shannon entropy scores for mechanism predictions on AAA domains that violate hierarchy definitions vs. those that do not.

While our categorization of AAA domains into mechanistic classes captures many AAA domains, it is not expected to be exhaustive. In other words, there are likely additional mechanisms that have not been accounted for in our five-class system and examples that fit into our mechanistic framework but are too evolutionarily divergent to be cleanly differentiated. We therefore sought to identify metrics that would flag mechanistic outliers. Toward this end, we first looked at the abundance of hierarchy violations within our mechanistic predictions across a subset of the proteome-derived domains from H. sapiens, S. pombe and E. coli (**Fig. 6B**). Hierarchy violations occurred in mechanistic predictions for roughly 20-40% of the proteome-derived hits, and with similar rates based on training with domain sequences, or entire ORFs.

Next, we wanted to assess how neatly the unknown proteins fall into the categories for which our logistic regression model was trained. For this, we used Shannon entropy, which gives measure of how evenly distributed the probabilities are for a given AAA domain sequence between the five classes. This can be conceptualized as a measure of ‘indecision’ for our model, ranging from zero to log2(n). In **Fig. 6C**, we compare the distributions of Shannon entropy values obtained for mechanism (n=5, log_2_[n] = 2.32) from the subset of predictions in which hierarchy was violated. The fact that Shannon entropy and hierarchy violation give results that are in general agreement supports the idea that both metrics are useful for identifying instances where ground truth may fall outside of the selection of annotations that the model is being forced to choose from. Furthermore, we find that, on average, average Shannon entropy is consistently higher within the hierarchy violations for open reading frames, consistent with the idea that focusing on the domain level increases the chance of in-distribution inference of mechanistic function.

## Discussion

In this work, we interrogate how sequence is related to molecular function in AAA protein through the development of CatFunc, a flexible and elegant tool for inference of molecular function for proteins. The AAA superfamily is an ideal test case for this, as it comprises a structurally conserved but mechanistically diverse family protein domains. Overall, we find that the information needed to predict mechanism of AAA domain-containing proteins is encoded within the AAA domain itself, with the inference performing marginally lower when focused on domain sequences than with full ORF context. While it is possible that context was leaked through the ESM embeddings from its original training on entire ORFs, we see reasonable clustering with traditional phylogenetic analysis as well, indicating that mechanistic character is truly encoded within the AAA domain.

As part of our analysis, it was critical to expand our dataset to ensure diversity and representation of domain sequences with different mechanistic labels. In phylogenetic analysis of both the domain and ORF level sequences, the literature sequences are well separated amongst the expanded data, except for the initiator class, whose representation is inflated due to its resistance to homology filtering. LD and phylogenetic analysis of the expanded dataset of ORFs reveals clear clusters of our mechanistic labels, providing important qualitative metrics for encoding of our mechanistic labels within the dataset.

Notably, our per-layer calibration experiments show that the information used for classification emerged relatively early in the language model, indicating that the classification arises from more general evolutionary trends rather than specific residue-residue interactions. However, later layers also demonstrated high levels of recall, despite our use of sequence averaged embeddings for classification. Thus, future work will focus on 1) implementing a more residue specific pipeline that can identify which residues give rise to classification of the model and 2) identifying examples where early and late layers of the model give different predictions and assessing this as a proxy for convergent vs. divergent evolution.

The cases where CatFunc misclassified domains during validation provide evolutionary insight into the development of the domain-encoded mechanistic labels, especially when cross-referenced with phylogenetic analysis. For example, the rotary motor domains misclassified as reciprocating motors appeared in an area where many initiators and rotary motors were phylogenetically outliers surrounded by reciprocating motors. Another phylogenetic cluster of misclassified rotary motors were called as initiators near a cluster of reciprocating motor domains, with additional rotary motors misclassified as reciprocating motors near the edge of this cluster. Taken with the fact that both the loader and signaling classes were commonly misclassified as rotary motors, these analyses are consistent with the idea that the rotary motor mechanism is the most ancient, and that the other mechanisms share an evolutionary lineage that traces back to it.

Last, we demonstrate the hierarchical classification is particularly useful for out-of-distribution functional inference, as it can be used to mark outliers from the functional schema being projected onto unknown examples. This metric, combined with the associated Shannon entropy values provide a confidence metric not just for the accuracy of a specific functional annotation, but whether a new, previously unobserved annotation is present within the sequence being queried. The most useful application of our pipeline is functional inference on AAA domains from out-of-distribution domain architectures; however, these require experimental validation that was beyond the scope of this work but will be pursued in future studies.

Approaching the sequence-function problem at a general level faces several challenges and is often focused on the prediction of functional annotation in the form of Gene Ontology terms. Here we demonstrate the advantage tailor-made functional questions to be formulated on a case-by-case basis, and that relatively simple PLM-based inference approaches can generate hypotheses that could potentially reduce the amount of time for an experimental research campaign to verify ground truth by months or years. To be effective, such approaches must be capable of approaching such mechanistic questions with user-defined nuance, we expect that future experimental campaigns will use inference-based approaches such as CatFunc to prioritize mechanistic outcomes based on examples from the literature. This framework will likely prove useful in the study of additionally biomedically important proteins such as kinases, membrane transporters and G protein-coupled receptors.

## Supplemental Figures

**Extended Data Figure 1:**
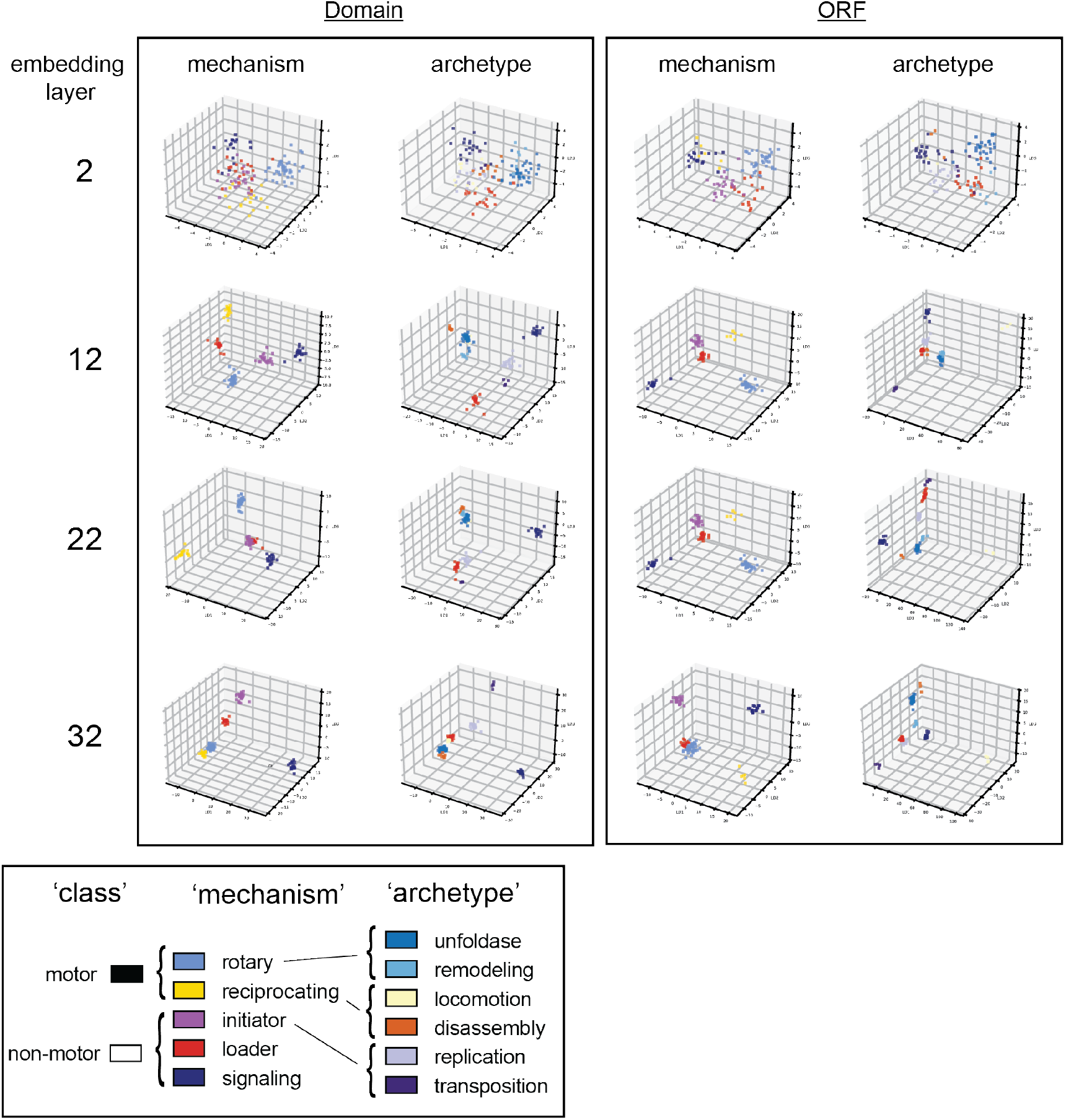
Selected ESM embedding layers of foundational AAA domains in categorical discriminant space. ESM2-derived protein sequence embeddings of AAA domains (left) or entire AAA domain containing ORFs from the foundational (literature) dataset projected into linear discriminant space for either mechanism or archetype labels. Actual labels from the dataset are mapped onto each sequences embeddings with the coloring scheme used throughout this manuscript.

**Extended Data Figure 2:**
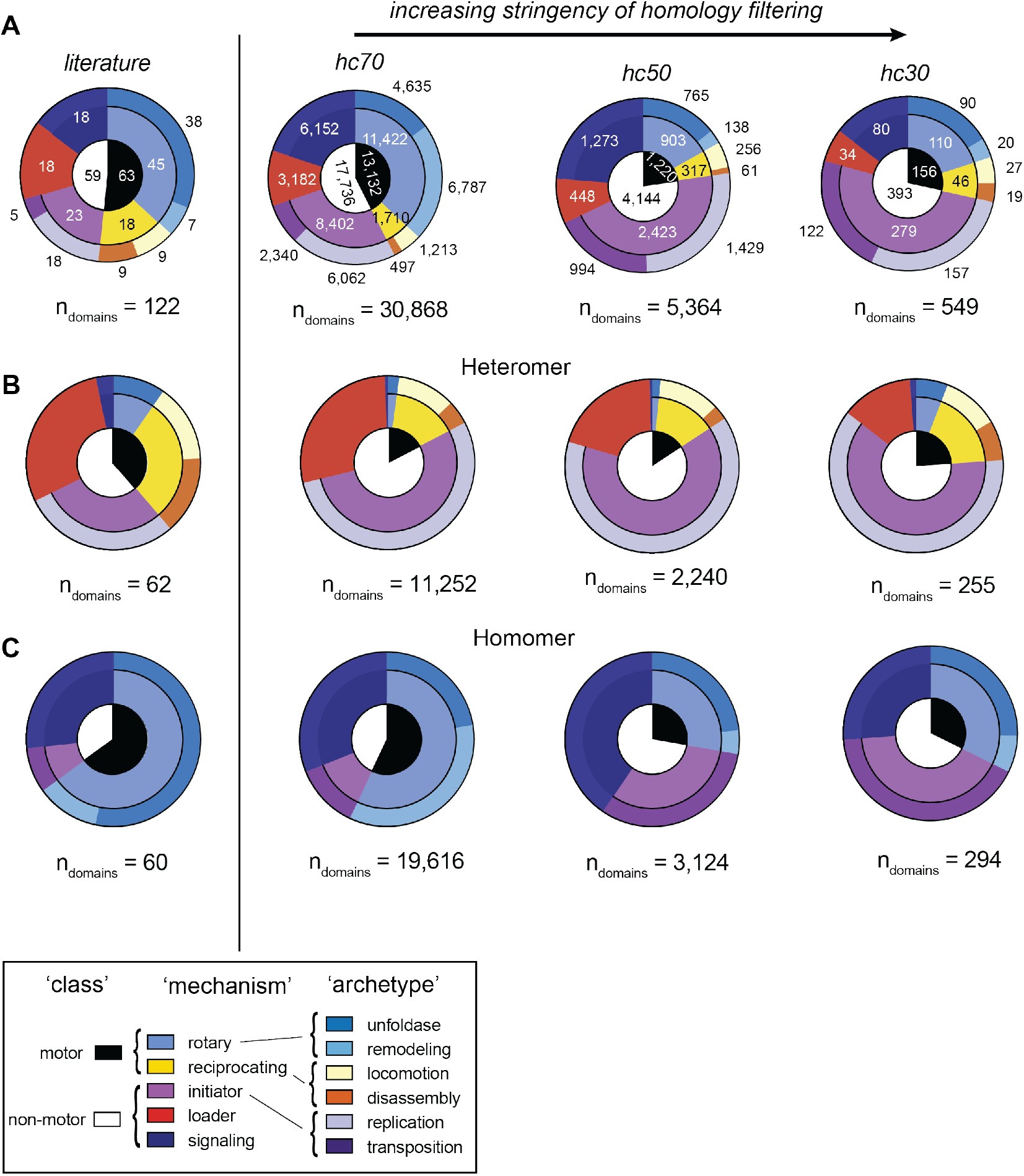
Label distribution as a function of homology stringency and oligomeric state in expanded dataset. **A**) Pie charts depicting the distribution and total number of hierarchical labels within the foundational (literature) dataset of AAA domains, and at 3 different homology cutoffs. Colored using the scheme used throughout this manuscript. **B**) Distributions of the same hierarchical labels within asymmetric ‘oligomeric state’ groups.

**Extended Data Figure 3:**
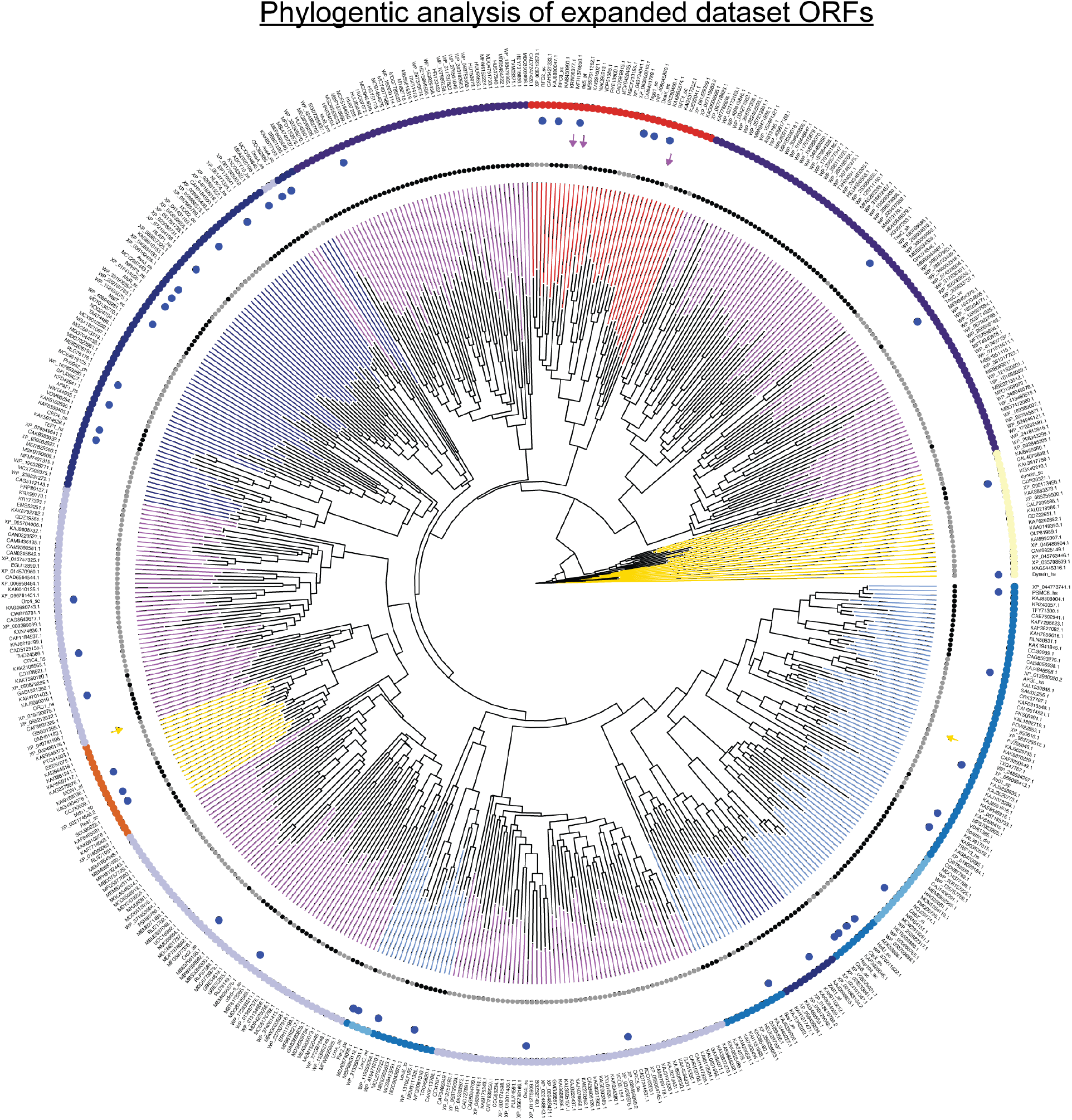
Phylogenetic analysis of ORFs from expanded dataset. Tree annotations are as in **Fig 5D**. Mechanistic labels are depicted as radial bars, validation group depicted as gray circles. misclassified results are colored arrows, colored according to the erroneous predicted mechanism. Blue circles denote the literature sequences, and outer radius depicts archetype labels using the coloring scheme used throughout this manuscript.

**Extended Data Figure 4:**
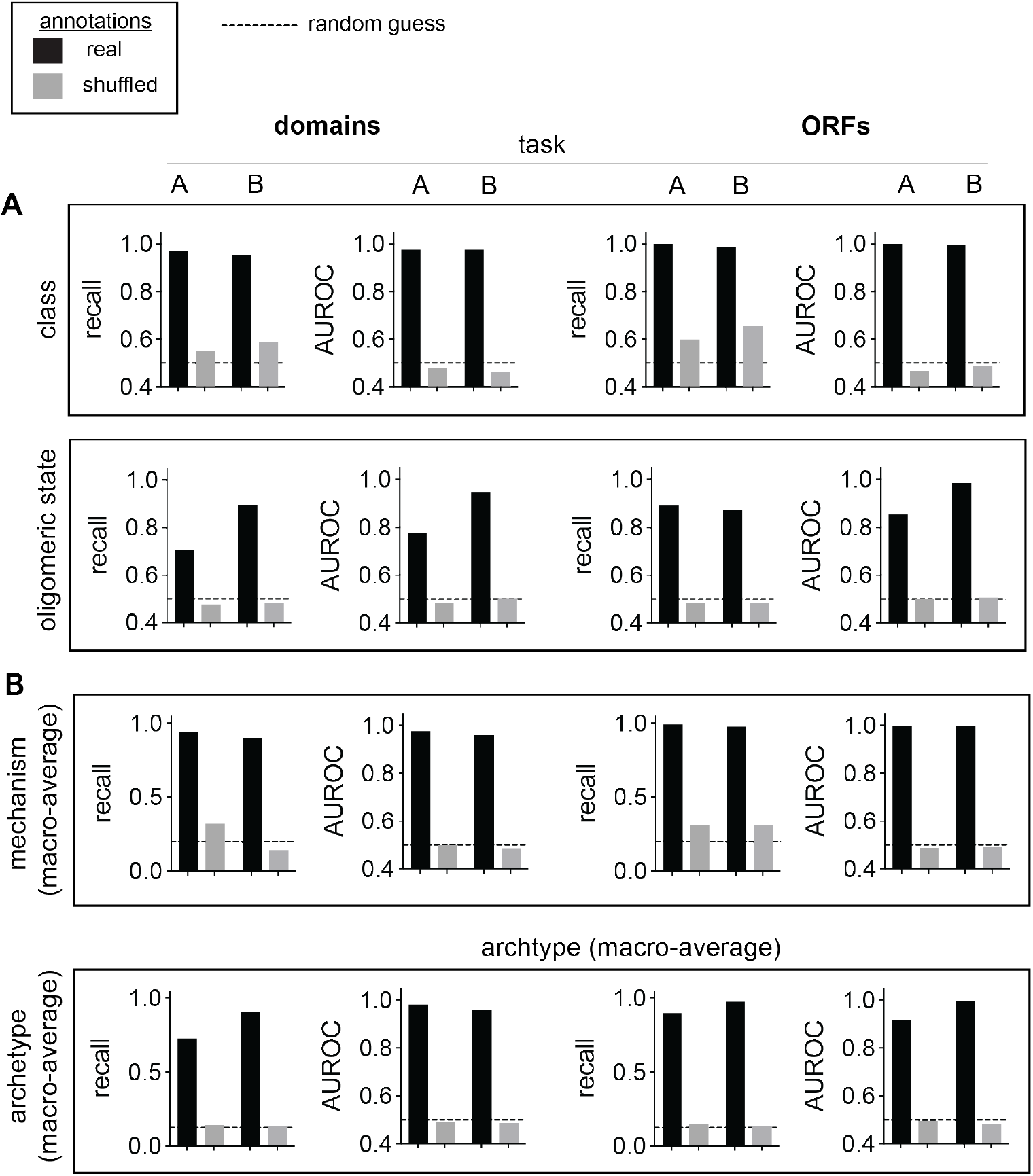
Extended binary and macro validation data. Recall and Area Under the Receiver-Operator Curve (AUROC) for **A**) the binary ‘class’ and oligomeric state labels. **B**) Macro-class-average recall and One vs Rest (OvR) AUROC for multi-class logistic regression in the two validation tasks using domain or ORF representations. Tasks A and B denote opposite usage of the two validation groups for training and testing of the logistic regression model. The model was trained on label-paired global ESM representations (layers 11-19, see Methods) of either domain (left) or ORF (right) sequences. Actual label-sequence associations were test alongside random, shuffled label associations. Dotted lines denote the values expected for a random guess.

**Extended Data Figure 5:**
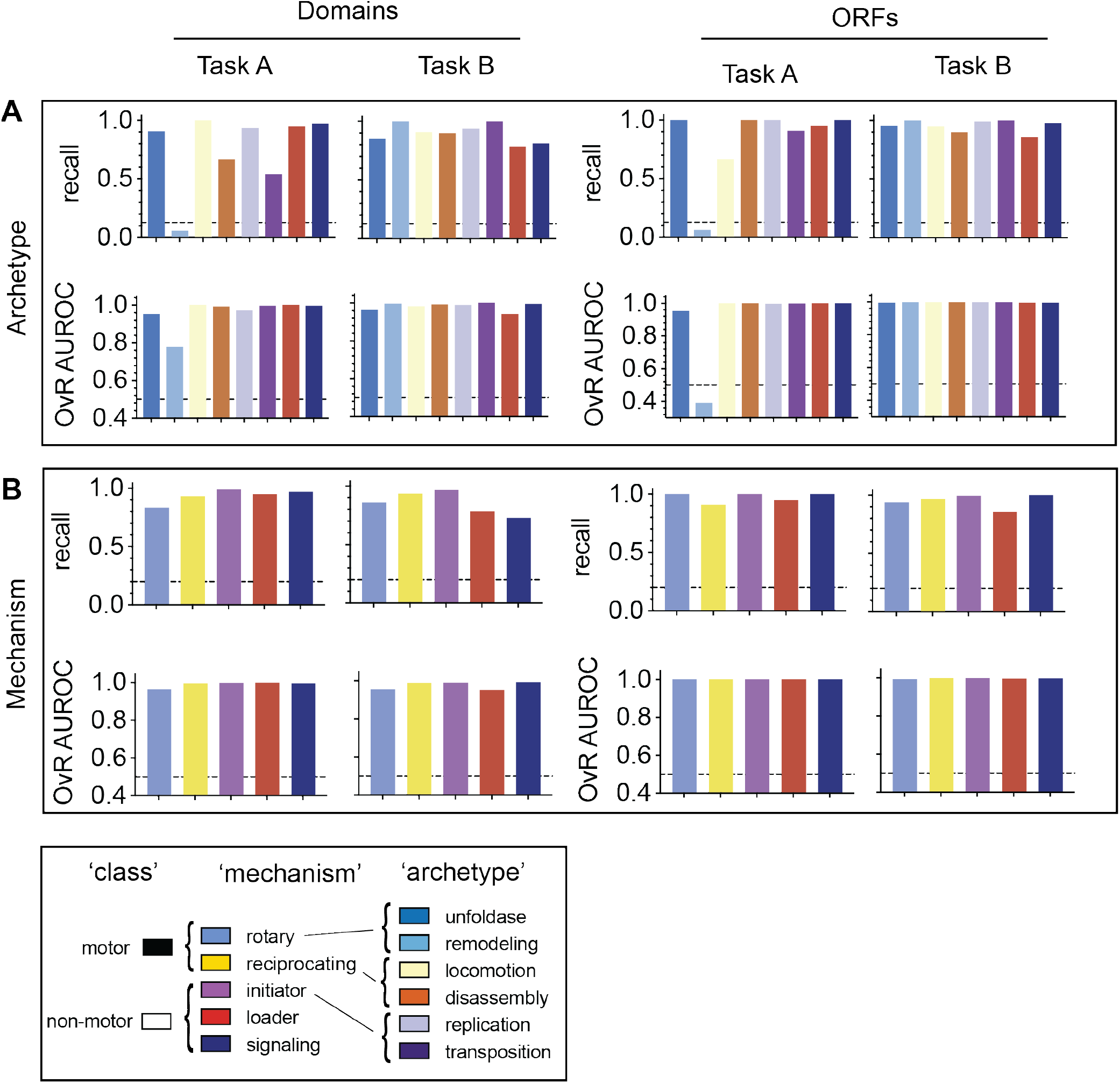
Extended per-class model performance data. Per-class recall and One-vs.-Rest (OvR) Area Under the Receiver Operator Curve (AUROC) data for **A**) archtype and **B**) mechanism labels of the expanded AAA dataset. The statistics for each label are depicted as bars bearing the color from the scheme used throughout this manuscript.

**Extended Data Figure 6:**
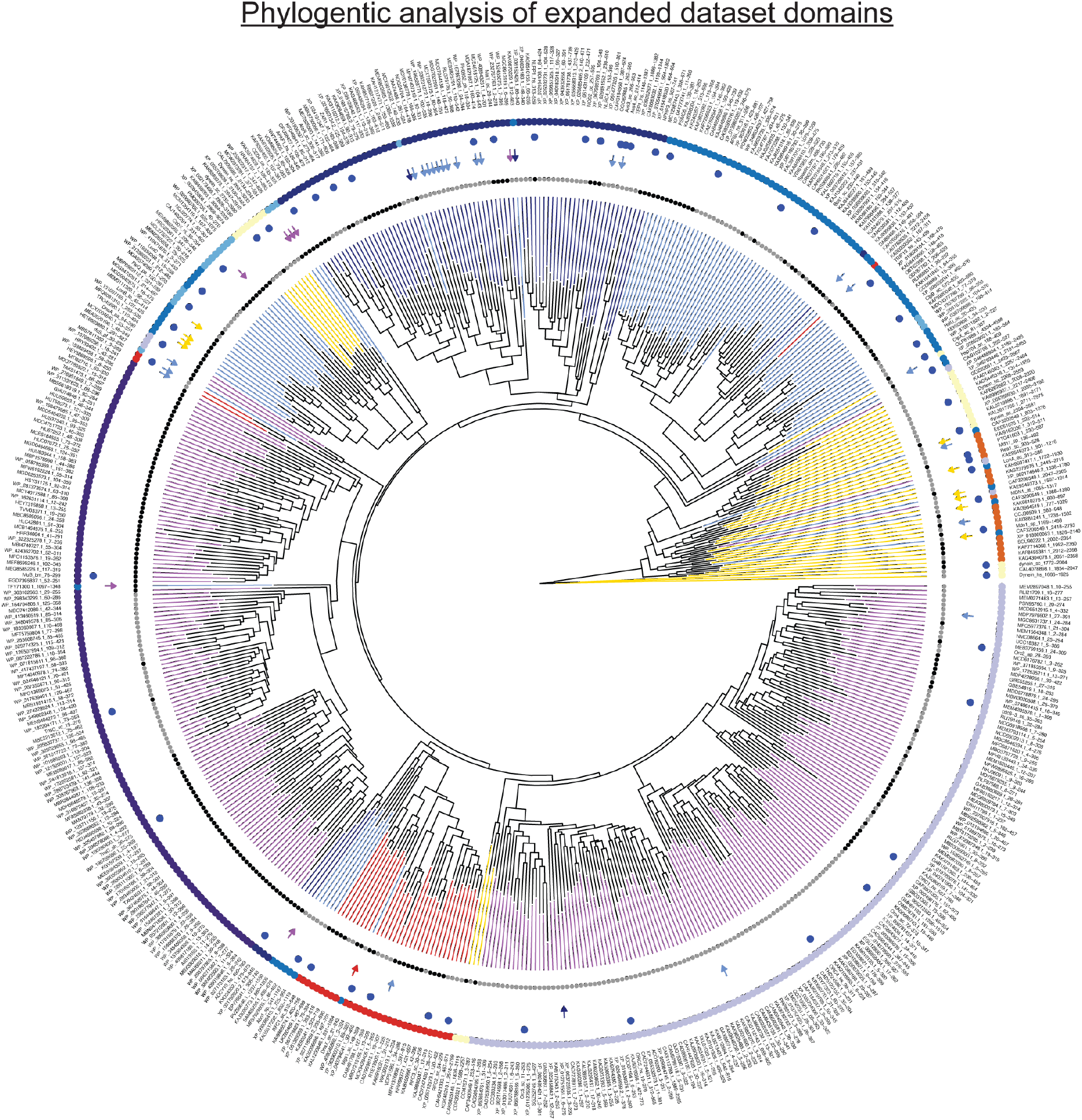
Phylogenetic analysis of domains from expanded dataset. Phylogenetic analysis of ORFs from expanded dataset. Tree annotations are as in **Fig 5D**. Mechanistic labels are depicted as radial bars, validation group depicted as gray circles. misclassified results are colored arrows, colored according to the erroneous predicted mechanism. Blue circles denote the literature sequences, and outer radius depicts archetype labels using the coloring scheme used throughout this manuscript.

## Supplementary Tables

**Extended Data Table 1:**
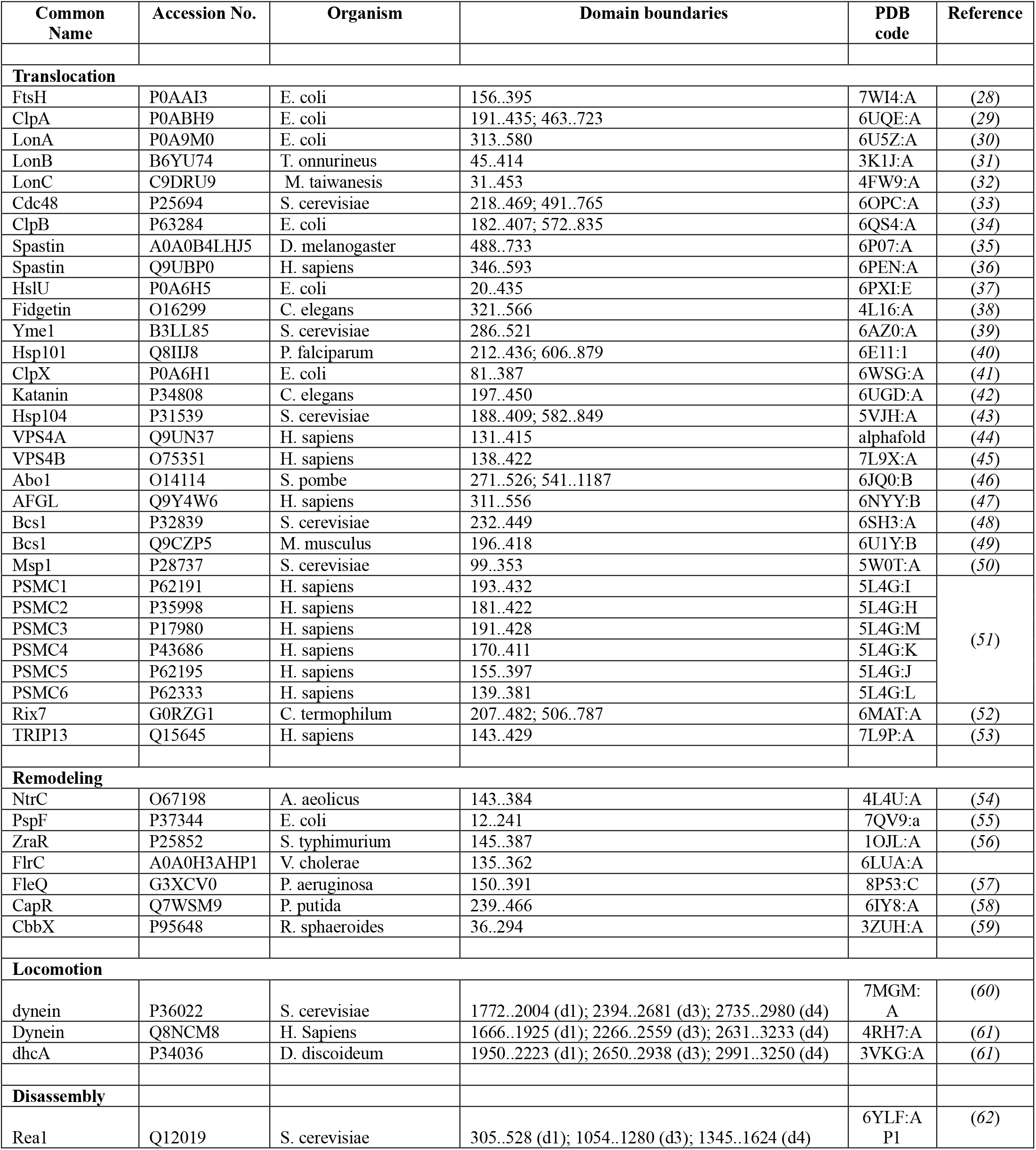
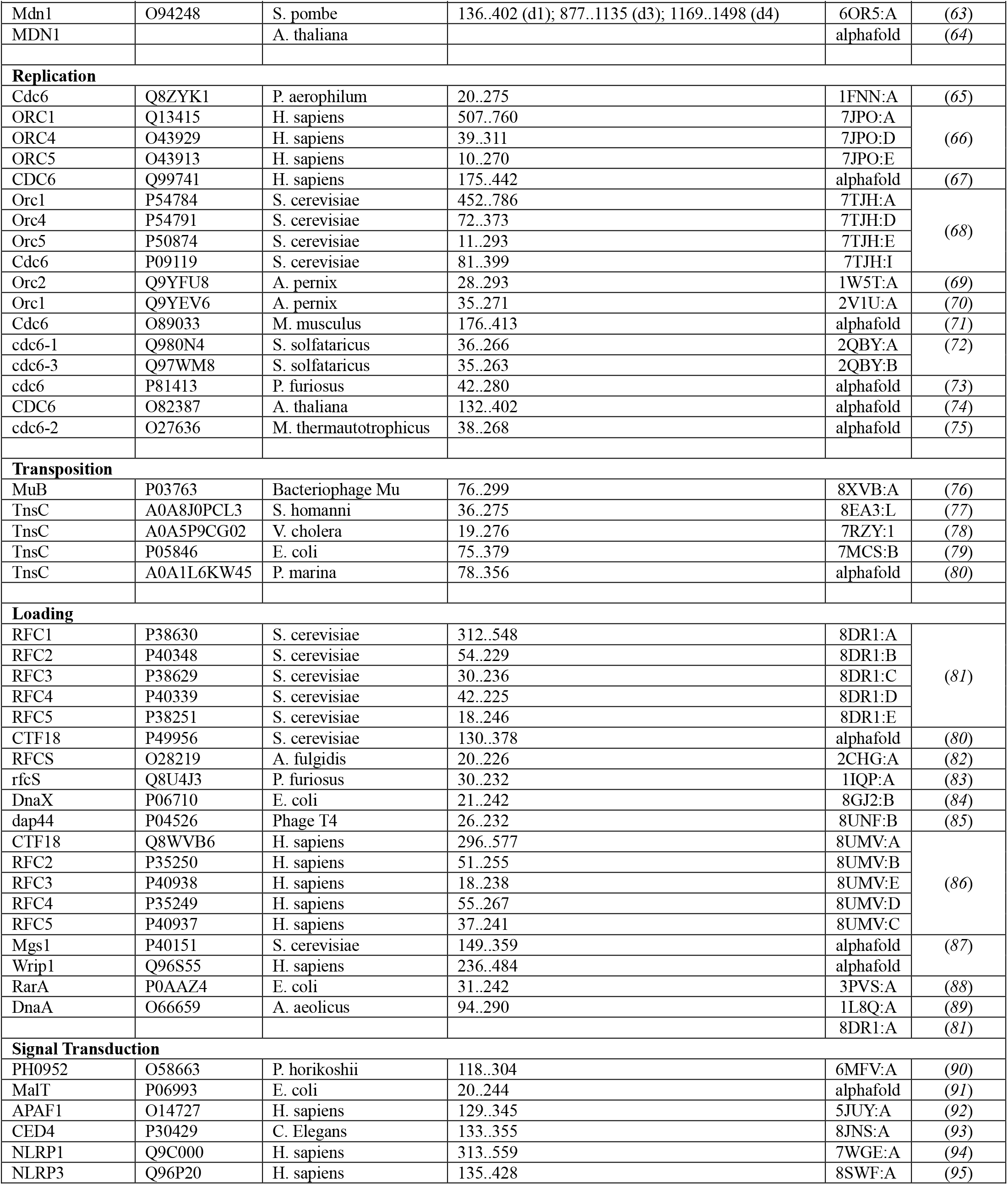
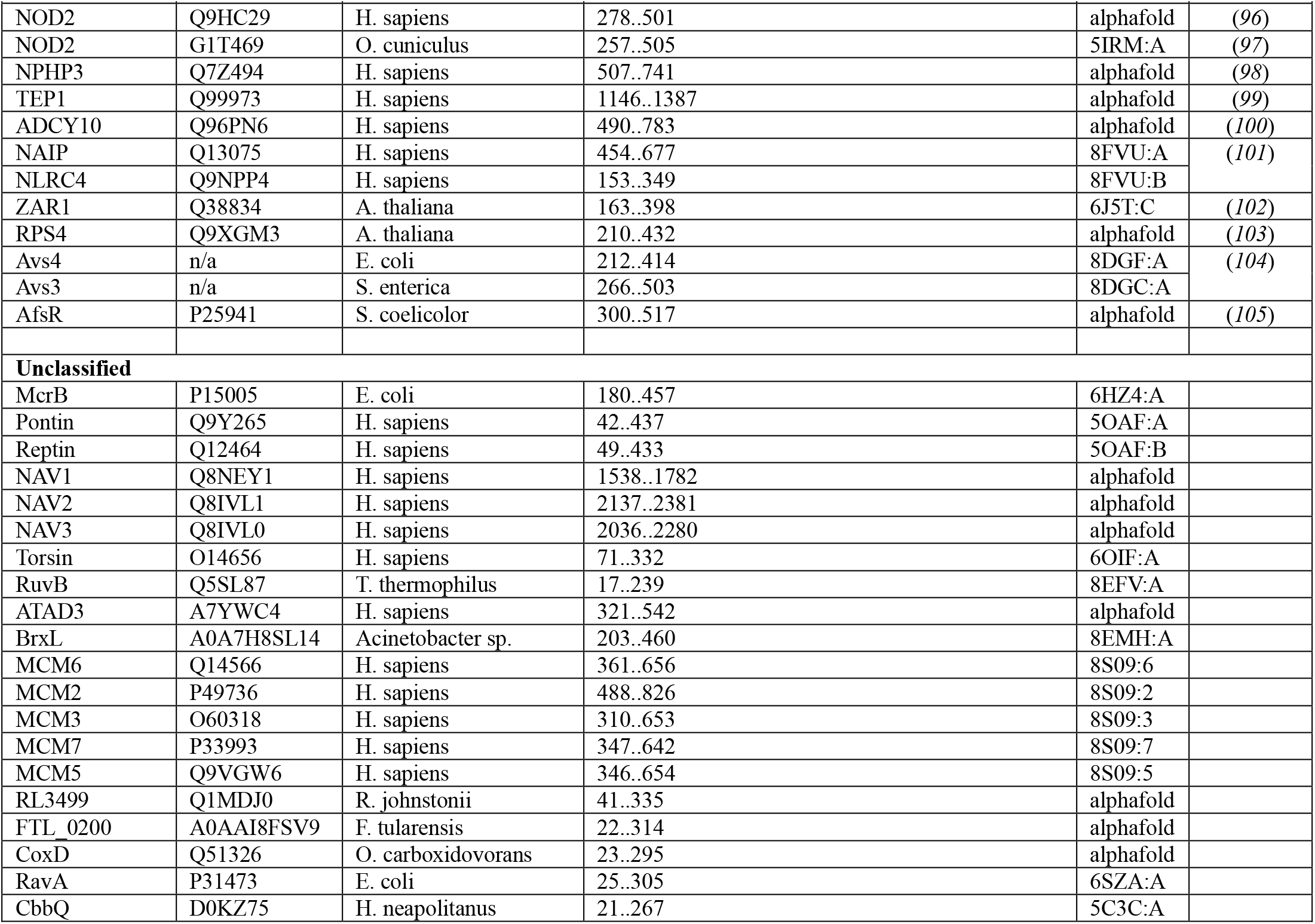
Foundational set of AAA domain-containing proteins from literature search.

**Extended Data Table 2:** Annotation prediction for selected AAA domains.

|  |  |  | domain |  |  |  | ORF |  |  |  |
| --- | --- | --- | --- | --- | --- | --- | --- | --- | --- | --- |
|  |  |  | mechanism |  | archetype |  | mechanism |  | archetype |  |
| description | accession | domain boundaries | prediction | entropy | prediction | entropy | prediction | entropy | prediction | entropy |
| McrB | P15005 | 180..457 | rotary | 1.12E-05 | unfoldase | 1.67E-05 | rotary | 1.106 | unfoldase | 0.939 |
| Pontin | Q9Y265 | 42..437 | rotary | 9.17E-01 | unfoldase | 5.94E-01 | rotary | 0.029 | unfoldase | 0.375 |
| Reptin | Q12464 | 49..433 | rotary | 1.00E+00 | unfoldase | 8.71E-01 | rotary | 0.017 | unfoldase | 0.397 |
| NAV1 | Q8NEY1 | 1538..1782 | recip | 7.41E-01 | locomotion | 6.95E-01 | initiator | 0.160 | replication | 0.291 |
| NAV2 | Q8IVL1 | 2137..2381 | recip | 5.68E-01 | locomotion | 3.39E-01 | initiator | 0.108 | replication | 0.191 |
| NAV3 | Q8IVL0 | 2036..2280 | rotary | 6.45E-01 | locomotion | 6.97E-01 | initiator | 0.096 | replication | 0.180 |
| Torsin | O14656 | 71..332 | recip | 6.76E-01 | unfoldase | 2.79E-01 | rotary | 0.011 | unfoldase | 0.020 |
| RuvB | Q5SL87 | 17..239 | recip | 6.76E-01 | loading | 1.33E+00 | rotary | 0.699 | replication | 0.953 |
| ATAD3 | A7YWC4 | 321..542 | rotary | 3.35E-04 | unfoldase | 2.13E-04 | rotary | 0.000 | unfoldase | 0.000 |
| BrxL | A0A7H8SL14 | 203..460 | rotary | 1.71E-03 | unfoldase | 6.40E-01 | rotary | 0.606 | unfoldase | 0.402 |
| MCM6 | Q14566 | 361..656 | recip | 6.29E-01 | unfoldase | 4.41E-01 | rotary | 0.074 | unfoldase | 0.065 |
| MCM2 | P49736 | 488..826 | rotary | 7.22E-01 | unfoldase | 2.90E-01 | rotary | 0.124 | unfoldase | 0.158 |
| MCM3 | O60318 | 310..653 | rotary | 5.84E-01 | unfoldase | 1.82E-01 | rotary | 0.490 | replication | 0.718 |
| MCM7 | P33993 | 347..642 | recip | 7.11E-01 | unfoldase | 4.09E-01 | initiator | 0.033 | unfoldase | 0.044 |
| MCM5 | Q9VGW6 | 346..654 | rotary | 7.26E-01 | unfoldase | 3.05E-01 | rotary | 0.053 | unfoldase | 0.061 |
| RL3499 | Q1MDJ0 | 41..335 | rotary | 6.72E-01 | unfoldase | 3.49E-01 | rotary | 0.379 | unfoldase | 0.381 |
| FTL_0200 | A0AAI8FSV9 | 22..314 | rotary | 2.82E-01 | unfoldase | 1.60E-01 | rotary | 0.516 | unfoldase | 0.450 |
| CoxD | Q51326 | 23..295 | rotary | 9.56E-02 | unfoldase | 3.79E-02 | rotary | 0.253 | unfoldase | 0.230 |
| RavA | P31473 | 25..305 | rotary | 2.56E-01 | unfoldase | 8.52E-01 | rotary | 1.026 | disassembly | 1.272 |
| CbbQ | D0KZ75 | 21..267 | rotary | 5.87E-01 | unfoldase | 4.57E-01 | rotary | 0.555 | unfoldase | 0.887 |

**Extended Data Table 3:**
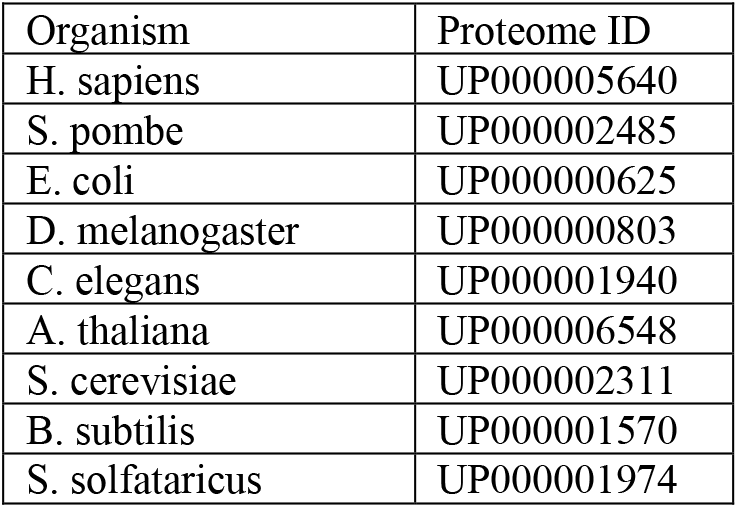
Proteome IDs used in HMMer scans for AAA domains.

## Supplemental Data

**Supplemental Data 1:** updated_literature.csv

**Supplemental Data 2:** lit_unk_50.aln

**Supplemental Data 3:** filtering_summary.csv

**Supplemental Data 4:** updated_expanded_dataset.csv

**Supplemental Data 5:** updated_unknowns.csv

**Supplemental Data 6:** survey_hits_keyword_false.csv

**Supplemental Data 7:** survey_hits_keyword_true.csv

## Methods

### Curation of AAA domain list

The complete list of foundational AAA domains is available as Table S1. To be included in this list, AAA domains with experimental structures were prioritized. Some exceptions were made in cases where there was sufficient functional data to support that they shared a mechanism with another member of the foundational set.

For the foundational list of AAA domains, boundaries were determined by manual inspection of experimental or predicted structures, with predicted structures obtained from the AlphaFold database in all cases except for Midasin from *A. thaliana*, where a structure was computed using the online AlphaFold server.

### Dimension reduction

Linear discriminant (LD) and principle component analysis (PCA) were implemented using their respective functions in scikit learn. Separate steps were used for fitting and projection in LD analysis to analyze the expanded dataset using discriminants derived from the literature set alone. PCA was performed using 95% variance preservation threshold. Scaling and PCA were done immediately prior to logistic regression, using the combined training and test embedding for that particular task. In cases where multiple embedding layers were pooled, dimension reduction was done subsequent to pooling.

### Dataset expansion

The initial dataset of AAA domain containing proteins was expanded using the Basic Local Alignment Search Tool (BLAST) (*106*), querying each open reading frame from the foundational set of AAA proteins against the NCBI non-redundant protein database, collecting 20,000 hits per query. Hits were then filtered based on coverage and homology score. Coverage was calculated as alignment length (a standard BLAST output) divided by the length of the query sequence. BLAST hits with coverage ratios between 0.8 and 1.1 and homology of at least 25% were considered sufficient quality to transfer categorical annotations from query.

To identify the boundaries of the AAA domains a hidden Markov profile was generated using HMMer (*107*) based on the foundational set of both annotated and unannotated AAA domains with a 50% pairwise homology maximum enforced (see bleow for details on homology filtering). The resulting profile was then applied to the filtered BLAST hits. The resulting domain sequences were then filtered with a maximum c-value of 1.0e-4 and sequence length between 190 and 500. Additionally, a custom scoring function was also used to filter based on walker A domain strength, identifying adjacent GK residues in the identified domains and scoring the upstream and downstream residues for similarity to the GxxGK(T/S) consensus walker sequence.

### Homology Filtering

All homology filtering in this project was executed using MMseqs2 (*108*) using the ‘easy-cluster’ command with cluster-mode of 1 and k = 5. The set of domains resulting from each individual BLAST query were homology filtered with a 95% pairwise maximum for deduplication. Subsequently, the datasets from the different queries were combined and filtered separately. Filtering was done in three sequential steps to achieve subset datasets at 70%, 50% and 30% pairwise homology maxima. For each homology level, representatives were chosen from the cluster such that underrepresented mechanistic labels would be prioritized. The label for prioritization is tunable in CatFunc. To ensure that the representative seqauences included the ground truth AAA domains from the literature search, the foundational set of AAA domains was filtered using the same parameters and logic. The datasets were then combined such that the foundational domains were included by combining the datasets for second round of clustering and removing the domains from the expanded dataset that clustered with member of the foundational set.

The most stringent, 30% pairwise homology maximum set of ORF was used for the downstream analysis, but the other datasets are included in Data S3 and Data S4. The expanded ORF dataset comprises the open reading frames from the domains that passed the final round of homology filtering.

### Inference of categorical annotations

Global sequence embeddings were generated using the ESM2 650M parameter model using the ‘mean’ option to average the embeddings across the protein sequences. For ORF sequence that exceeded the 1022 amino acid limit for represention by ESM2, the sequences were split into 1022 residue chunks with 150 amino acids worth of overlap between adjacent chunks. The final chunk was also 1022 residues long, but had additional overlap with the penultimate chunk to accommodate the remainder. Global embeddings of each chunk were then averaged to generate an embedding for the full ORF.

The scikit-learn Python package was used for all downstream machine learning tasks. Preprocessing was performed using the StandardScaler and PCA functions, with PCA set to retain 95% variance. Preprocessing was performed just upstream of model training on the set of sequences to be tested. In experiments where multiple layers were combined, Preprocessing was performed subsequently to pooling the layered embeddings. Supervised classification was performed using scikit-learn’s logistic regression implementation, with the regularization parameter, *C*, set to 0.1.

### Phylogenetic analysis

Multiple sequence alignments were performed using MAFFT (*109*), using Needleman-Wunsch (global pairing) for domain sequences and Smith-Waterman (local pairing) for full ORFs. Example commands are given below.

mafft --maxiterate 1000 --globalpair --retree 3 [*domains.fasta*] > [*aligned_domains.aln*]

mafft --maxiterate 1000 --localpair --retree 3 [*orfs.fasta*] > [*aligned_orfs.aln*]

Trees were then calculated using IQ-TREE (v3.0.1) (*110*): and were visualized with a custom R script using the phytools (*111*) and treeio (*112*) packages to map mechanistic annotations onto phylogenetic trees.

### Proteomic surveys

Proteome surveys were conducted using Hidden Markov scans on the ‘one-protein-per-gene’ proteomes available at UniProt. Scans were performed using HMMer, using the same profile and filtering criteria that was generated for dataset expansion. Upon inspection of the proteins for which HMMer have a positive signal, many had annotations as ABC-transporters. Hits with annotations for ABC transporters were removed by keyword search against the annotation text.

## Data and code availability

All data for these studies are available in the supplementary data files. All code is available at https://github.com/SwanStar

## Acknowledgements

J.A.S. would like to thank the Jane Coffin Childs foundation for funding this research. We thank Tyler Starr, Hannah Wayment-Steele and Assaf Elazar for key scientific input on this work. We thank Chris Miles, Aaron Boakye, Heidi Schubert and Tyler Starr for feeddback on the manuscript. The support and resources from the Center for High Performance Computing at the University of Utah are gratefully acknowledged.

